# Differential attenuation of β2 integrin-dependent and -independent neutrophil migration by Ly6G ligation

**DOI:** 10.1101/458760

**Authors:** Pierre Cunin, Pui Y. Lee, Edy Kim, Angela B. Schmider, Nathalie Cloutier, Alexandre Pare, Matthias Gunzer, Roy J. Soberman, Steve Lacroix, Eric Boilard, Craig T. Lefort, Peter A. Nigrovic

## Abstract

Abstract

Antibody ligation of the murine neutrophil surface protein Ly6G disrupts neutrophil migration in some contexts but not others. We tested whether this variability reflected divergent dependence of neutrophil migration on β2 integrins, adhesion molecules that interact with Ly6G at the neutrophil surface. In integrin-dependent murine arthritis, Ly6G ligation attenuated joint inflammation, even though mice lacking Ly6G altogether developed arthritis normally. By contrast, Ly6G ligation had no impact on integrin-independent neutrophil migration into inflamed lung. In peritoneum, the role of β2 integrins varied with stimulus, proving dispensable for neutrophil entry in *E. coli* peritonitis but contributory in IL-1-mediated sterile peritonitis. Correspondingly, Ly6G ligation attenuated only IL-1 peritonitis, disrupting the molecular association between integrins and Ly6G and inducing cell-intrinsic blockade restricted to integrin-dependent migration. Consistent with this observation, Ly6G ligation impaired integrin-mediated postadhesion strengthening for neutrophils arresting on activated cremaster endothelium *in vivo*. Together, these findings identify selective inhibition of integrin-mediated neutrophil emigration through Ly6G ligation, highlighting the marked site and stimulus specificity of β2 integrin dependence in neutrophil migration.

**KEY POINTS:**

- The contribution of β2 integrins to neutrophil migration into inflamed tissues varies with site and stimulus.
- Ligation of Ly6G, a GPI-linked neutrophil surface protein, selectively attenuates β2 integrin-dependent neutrophil migration in vivo.
- Blockade correlates with disrupted interaction between Ly6G and β2 integrins and impaired integrin-mediated postadhesion strengthening.

## Introduction

The entry of neutrophils into tissues enables immune defense but also pathologic inflammation. Egress from the vasculature is thus tightly controlled. Rolling is followed by arrest at the endothelial surface, whereupon neutrophils crawl toward sites favorable for diapedesis between or through endothelial cells. Neutrophil β2 integrins, in particular CD11a/CD18 (LFA-1) and CD11b/CD18 (Mac-1), play key roles in this process, mediating slow rolling, arrest, postadhesion strengthening, crawling, and migration across the endothelium.^1-3^ In patients lacking β2 integrins, defective neutrophil migration is believed to contribute to susceptibility to invasive bacterial infections.^4^

Yet neutrophil migration does not always require β2 integrins. Patients deficient in CD18 can still develop neutrophilic lung infiltrates.^5^ Correspondingly, experimental CD18 blockade does not impair neutrophil migration toward Streptococcus pneumoniae or C5a instilled into rabbit lung, although the response to IL-1α is impaired, and partial impairment is reported with *Escherichia coli* or *Staphylococcus aureus*.^6-8^ In murine lung, the impact of integrin blockade, deficiency, or manipulation varies strikingly with experimental conditions, in some contexts even enhancing neutrophil infiltration though facilitated release from adhesive attachments.^9-13^ Neutrophil recruitment independent of β2 integrins is observed in murine liver.^14^ Thus, β2 integrins can mediate neutrophil migration into tissues, but dependence on these molecules displays substantial variability.

The pathways mediating neutrophil egress have therapeutic implications. Clinical experience in neutropenic patients highlights the critical role of this lineage in immune defense. However, if neutrophils recruitment could be antagonized selectively, particularly in sterile inflammation, then it could be possible to drive a wedge between the pathogenic and protective capacities of neutrophils.

Several years ago, we observed that antibodies against the murine neutrophil surface protein Ly6G abrogates murine inflammatory arthritis, even without depletion of neutrophils from the circulation.^15^ This effect reflected blockade of neutrophil entry into the inflamed joint, and to a lesser degree into thioglycolate-stimulated peritoneum, correlating with interruption of migration in a β2 integrin-dependent system *in vitro*. Ly6G was found to interact at a molecular level with surface β2 integrins, suggesting a regulatory interaction between these molecules.^15^ Yet observations seemingly discordant with this possibility have been reported. The anti-Ly6G/Ly6C antibody RB6- 8C5 (also called anti-Gr-1) has been used in vivo to label neutrophils to study their migration in several tissues, including cremaster muscle, skin, and liver.^14,16-18^ While these studies did not compare antibody with control, RB6-8C5-treated mice resembled untouched animals with respect to the migration of fluorescent neutrophils to intradermal S. aureus.^19^ Neutrophils from Ly6G-deficient “Catchup” mice migrate normally.^20^ Thus, there remains substantial uncertainty about the functional capacity of this highly-abundant neutrophil marker.

We considered the possibility that the variable effect of Ly6G ligation reflected the variable contribution of β2 integrins to neutrophil migration. Using WT, CD18^-/-^ and Ly6G^-/-^ mice, we confirm that β2 integrin dependence varies strikingly with tissue and inflammatory stimulus. We find that Ly6G ligation selectively attenuates β2 integrin-driven neutrophil migration, potentially via a direct impact on the association of Ly6G with these integrins. This result suggests that targeting integrin-associated molecules such as Ly6G could exploit variable integrin dependence for selective targeting of neutrophil migration in some contexts while preserving their role in others.

## Results

### Ly6G ligation, but not Ly6G deficiency, attenuates K/BxN serum transfer arthritis

Ly6G ligation blocked neutrophil migration in IgG immune complex-mediated K/BxN serum transfer arthritis, yet Ly6G-deficient neutrophils exhibit no overt migratory defect.^15,20^ To reconcile these findings, we tested the effect of Ly6G deficiency in this neutrophil- and β2 integrin-dependent arthritis model.^21,22^ Littermate WT (Ly6G+/+) and Catchup mice (Ly6G+/- and Ly6G-/-) were treated with K/BxN serum 150μl i.p on day 0 and day 2 and arthritis was evaluated over 1 week as described.^15^ Ly6G deficiency had no impact on arthritis, as assessed by clinical score and joint thickening (Figure 1A and B). We then treated Ly6G^-/-^ and WT mice with the anti-Ly6G antibody 1A8 5μg i.p., a dose that does not reduce circulating neutrophil count, or the corresponding isotype control 2A3 at day −1 (one day before arthritis induction) and then again at day 2.^15^ Arthritis was abrogated by 1A8 in WT mice, whereas Catchup animals developed arthritis normally (Figure 1C and D). Together with previously published findings15, these data confirm that Ly6G ligation modulates neutrophil migration in this integrin-dependent model, and highlight the distinction between ligation and genetic deletion with respect to Ly6G.^20^

**Figure 1.**
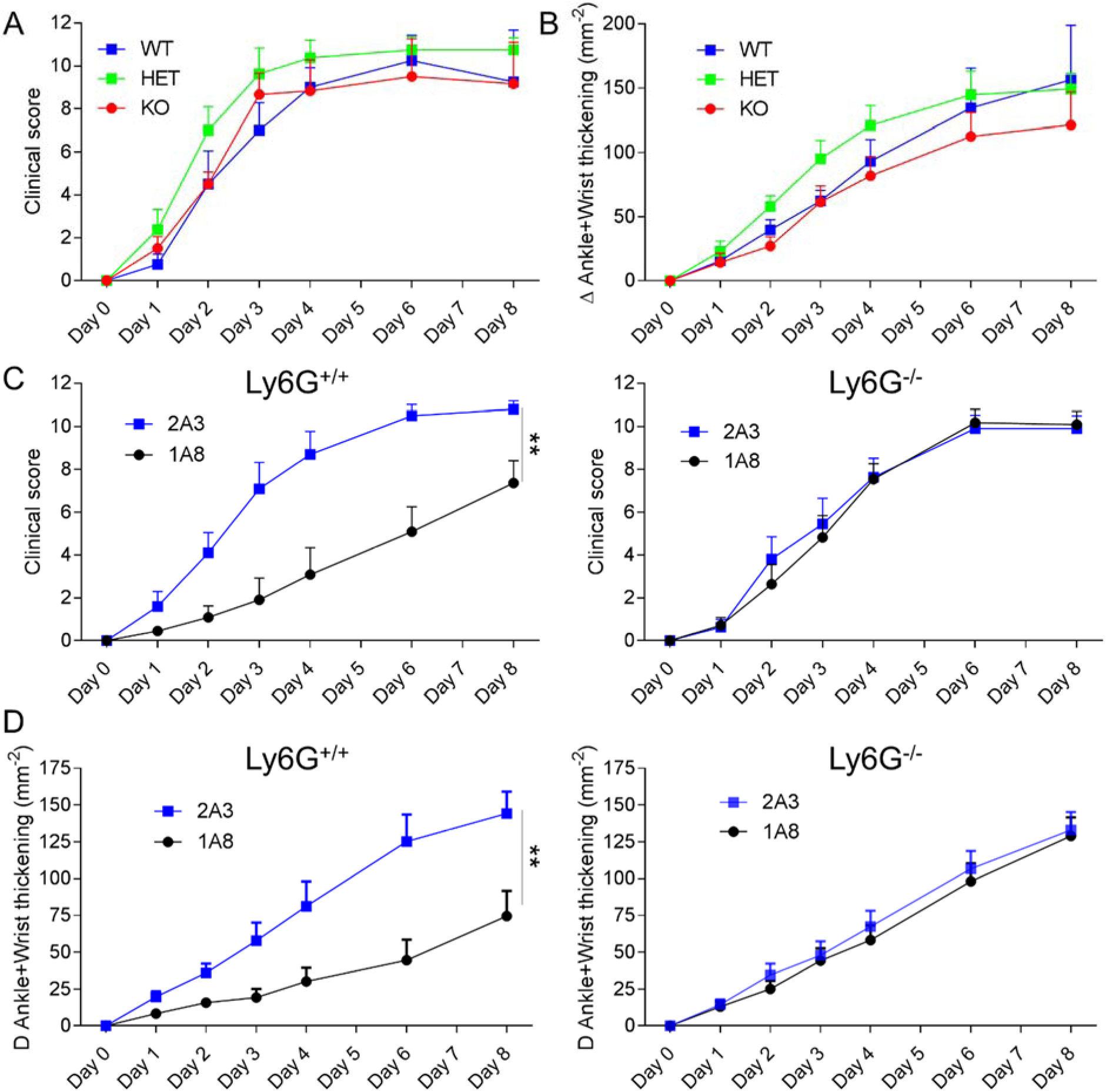
Ly6G ligation but not deficiency attenuates IgG-mediated arthritis. **A-B**. Male mice were treated with K/BxN serum 150μl i.p. on days 0 and 2 and assessed for arthritis over 8 days, measuring clinical scoring on a 0-12 scale (**A**) and change in ankle and wrist thickness (**B**). Results pooled from 2 identical experiments using a single pooled batch of K/BxN serum. Ly6G^+/+^ (WT, blue line, n=4), Ly6G^+/-^ (HET, green line, n=8) and Ly6G^-/-^ (KO, red line, n=6). Clinical scoring and thickness change: p=ns. **C-D**. WT and KO mice were treated with 5μg of 2A3 or 1A8 one day prior (day-1) and 2 days after (day+2) arthritis induction. **C**. Clinical scoring: 2A3- vs. 1A8-treated WT: p=0.0013; 2A3- vs. 1A8-treated KO: p=ns. **D**. Thickness change: 2A3- vs. 1A8-treated WT: p=0.0058; 2A3- vs. 1A8-treated KO: p=ns, n=10-11 mice / group.

### Ly6G ligation does not block integrin-independent neutrophil migration into lung

We then tested pneumonitis, in which the role of integrins in neutrophil recruitment diverges widely among published reports. To define β2 integrin dependence in our own setting, we transferred a 1:1 mixture of WT neutrophils stained with CellVue maroon (far-red) and CD18^-/-^ neutrophils stained with PKH67 (green) i.v. into mice previously treated i.n. with IL-1 β (25ng) or LPS (1μg). After 6 hours, we quantitated far-red and green neutrophils migrating into either lung parenchyma or airway (Figure 2A). WT and CD18^-/-^ neutrophils emigrated comparably to either stimulus, demonstrating that neutrophil migration under these conditions exhibits no cell-intrinsic requirement for β2 integrins; a subtle but statistically significant skew in favor of WT neutrophils (WT / CD18^-/-^ ratio ~1.28-1.41, p<0.05) was identified for neutrophils entering parenchyma but not airway (Figure 2B). We then studied the effect of Ly6G blockade. Mice were treated with 1A8 or the corresponding isotype control 2A3, as above. After 18 hours, mice received i.n. IL-1β to induce neutrophil migration into the lung, and neutrophils were counted in the airway and parenchyma 6 hours later (Figure 2C). Paralleling lack of integrin dependence, no effect of 1A8 on neutrophil entry was observed (Figure 2C). This result confirms that Ly6G ligation does not induce generalized impairment of neutrophil migration, either functionally or through effects on the quantity or quality of circulating neutrophils. Together with the arthritis data, these findings are consistent with (but do not prove) a selective effect of Ly6G ligation on β2 integrin-dependent migration.

**Figure 2.**
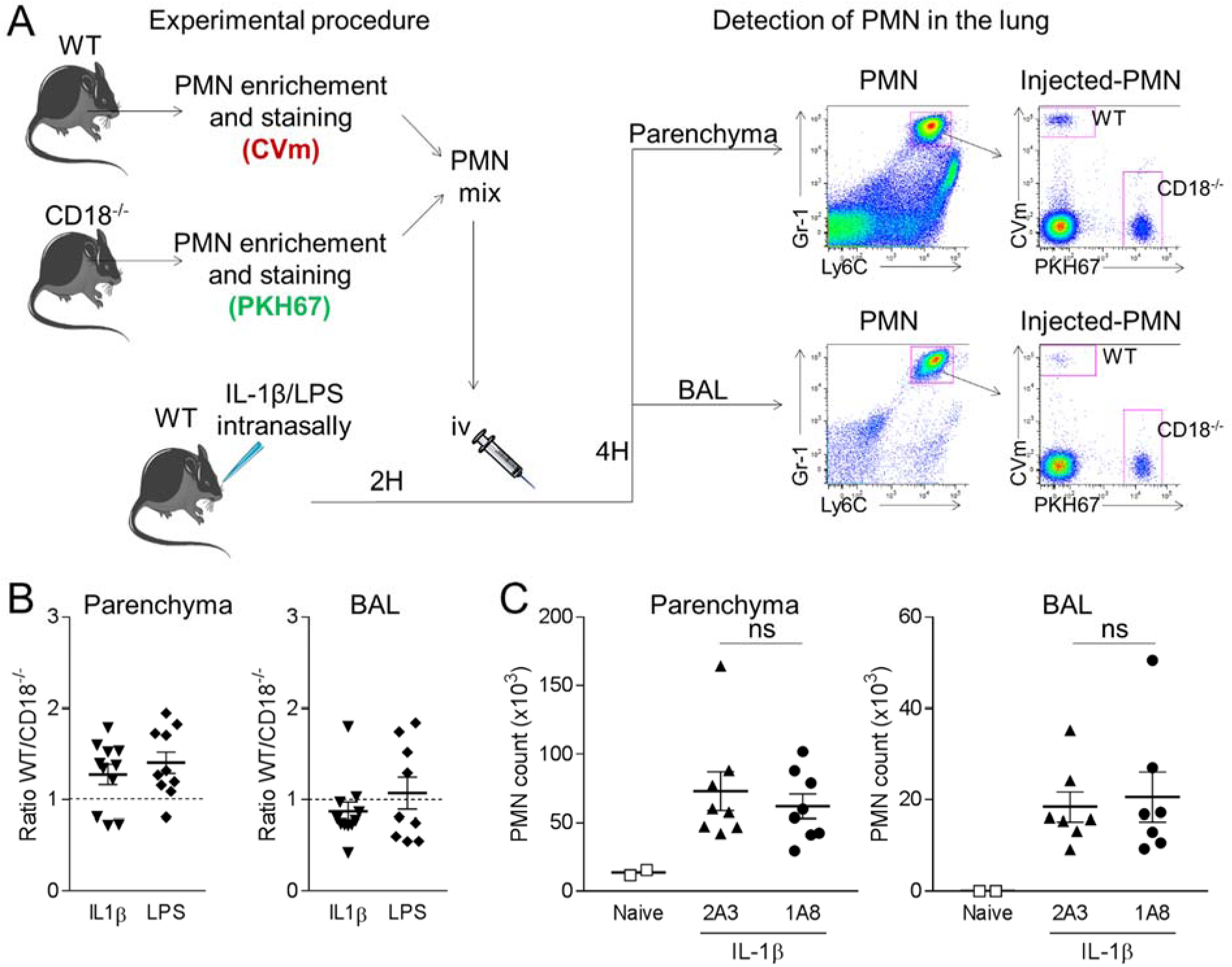
Neutrophil recruitment to lung does not require β2 integrins and is unimpaired by Ly6G ligation. **A**. PMN were enriched from WT and CD18^-/-^ marrow, stained with CellVue maroon (CVm, far-red) or PKH67 (green), and mixed at a 1:1 ratio (colors for WT and CD18^-/-^ cells varied across experiments). Cells were injected i.v. into mice treated intranasally with IL-1β (25ng) or LPS (1 μg) 2 hours previously. Four hours after cell injection, the presence of far-red PMN (WT) and green PMN (CD18-/-) in the bronchoalveolar lavage (BAL) and in lung parenchyma was assessed by FACS. **B**. Ratio WT/CD18-/- PMN in parenchyma and BAL fluid, n=9-11 mice per group; mean +/- SEM and deviation from 1:1 are: parenchyma IL-1β 1.28 +/- 0.11 p = 0.03, parenchyma LPS 1.41 +/- 0.12 p = 0.007; BAL IL-1β 0.87 +/- 0.10 p = ns, BAL 1.07 +/- 0.18 p = ns. **C**. Mice were injected i.p. with 5μg of 1A8 or 2A3 and 18 hours later 25ng IL-1β was administrated intranasally. PMN in the BAL and in lung parenchyma were counted by flow cytometry 6 hours later, n=7-8 mice per group.

### Neutrophil migration into murine peritoneum exhibits β2 integrin dependence that varies with inflammatory stimulus

Studies of the role of β2 integrins in murine peritonitis describe widely varying results, ranging from little role to marked dependence.^12,23-25^ We considered the possibility that this variability reflected differences in inciting stimulus. We therefore compared WT and CD18^-/-^ mice in sterile peritonitis mediated by IL-1β (5ng i.p. x 1) and septic peritonitis mediated by live E. coli (107 CFU i.p. × 1). These stimuli elicited robust neutrophil accumulation in both strains (Figure 3A). Interpretation of these findings was complicated by the circulating neutrophilia of CD18^-/-^ mice (^10,23,26,27^ and Figure 3B and C), and by the presence of an appreciable population of neutrophils even in unmanipulated CD18^-/-^ peritoneum (Figure 3C). Calculation of the ratio of migrating to circulating neutrophils suggested that β2 integrin deficiency impaired migration more in IL-1β than *E. coli* peritonitis, but an unambiguous difference was difficult to establish (Figure 3D).

To overcome this obstacle, we employed adoptive transfer as per our lung studies (Figure 3E). WT and CD18^-/-^ neutrophils were stained in green (PKH67) and far-red (CellVue maroon), respectively. These populations were mixed at a 1:1 ratio and injected i.v. into WT mice 1h after i.p. *E. coli* or IL-1β. At 3h, peritoneum was lavaged and relative migration of WT and CD18^-/-^ neutrophils enumerated by flow cytometry (Figure 3E). As suggested by the experiments in each strain alone, the role of β2 integrins was much more pronounced in IL-1β peritonitis (WT / CD18^-/-^ ratio 6.52 +/- 0.99, p<0.0001) than in *E. coli* peritonitis (WT / CD18^-/-^ ratio 1.91 +/- 0.20, p<0.001) (Figure 3F), although neutrophil migration in *E. coli* peritonitis remained partially β2-integrin dependent, with a ratio of migrating WT / CD18^-/-^ neutrophils approaching 2. Thus, consistent with the divergence among published results, the role of β2 integrins in neutrophil migration to the peritoneum varies markedly with stimulus.

**Figure 3.**
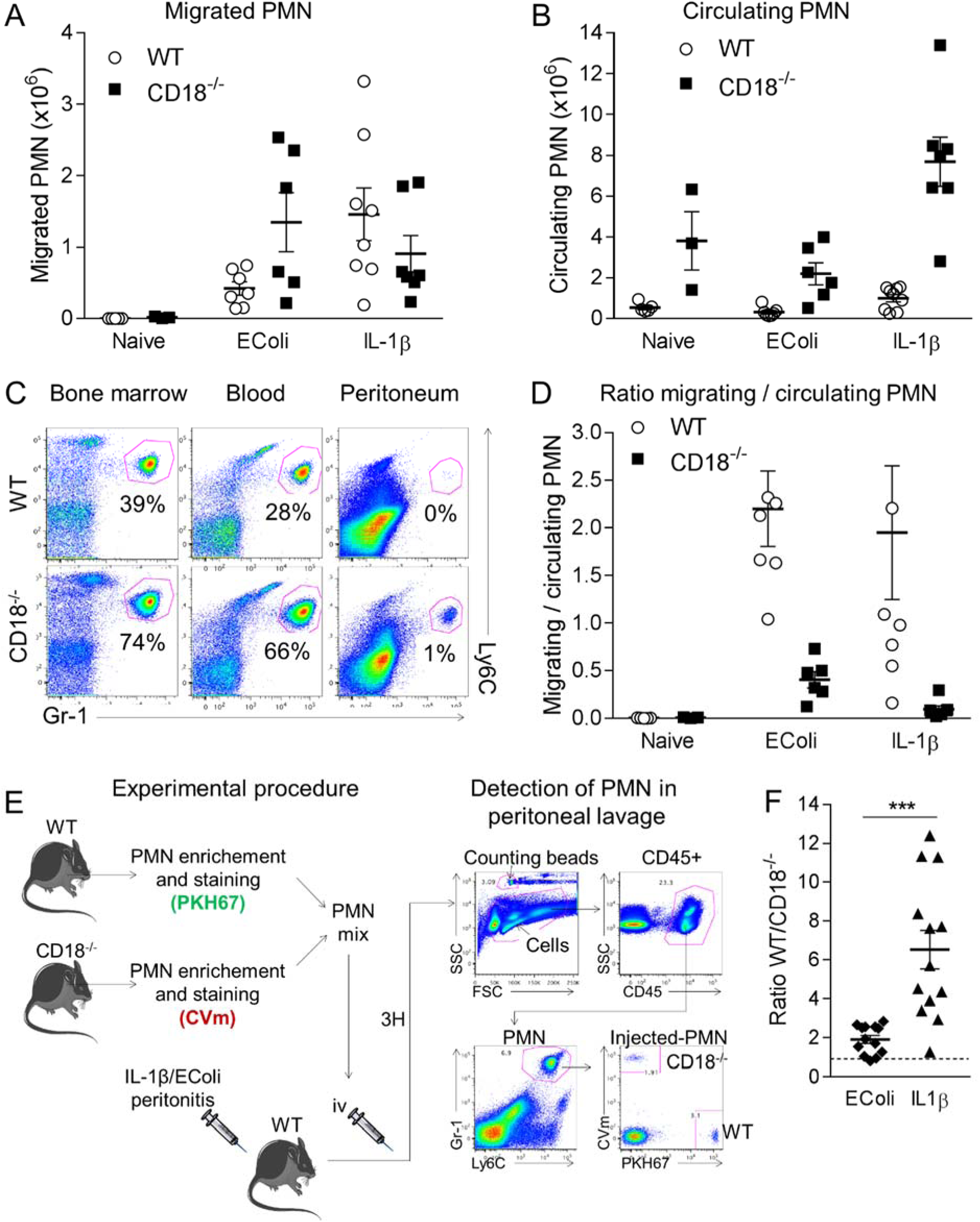
The dependence of neutrophils on β2 integrins for migration to peritoneum varies with stimulus. **A-B**. WT and CD18^-/-^ mice were injected i.p. with *E. coli* (10^7^ CFU/mouse) or IL-1β (5ng/mouse). Three hours later PMN in peritoneum lavage (**A**) and blood (**B**) were counted by FACS using counting beads (left histograms). **C**. Percentage of Gr-1/Ly6C-high PMN among CD45+ cells in WT and CD18^-/-^ bone marrow (left), blood (middle) and peritoneum lavage (right) was evaluated by FACS. Representative of at least 5 mice. **D**. Ratio migrated / circulating PMN, allowing an estimation of the relative PMN migration in WT and CD18^-/-^ mice, n=3-8 mice per group. **E**. PMN were enriched from WT and CD18^-/-^ marrow, stained with PKH67 (green) or CellVue maroon (CVm, far-red), and mixed at a 1:1 ratio (colors for WT and CD18^-/-^ cells varied across experiments). Cells were injected i.v. into mice treated with *E. coli* or IL-1 β i.p. 1 hour previously. Three hours after cell injection, the presence of green PMN (WT) and far-red PMN (CD18-/-) in peritoneal lavage was assessed by FACS. **F**. Calculation of the ratio of migrated WT / CD18^-/-^ PMN allows an estimation of the β2 integrin dependency in these 2 models of peritonitis, n=13 mice per group.

### Ly6G ligation specifically inhibits β2 integrin-dependent neutrophil migration

This variability in integrin dependence provided us an opportunity to test whether Ly6G ligation selectively inhibits β2 integrin-mediated neutrophil migration. Mice were treated with 1A8 or 2A3 as above, followed 3 days later by i.p. injection of either IL-1β or E. coli, a timepoint at which 1A8 remains detectable on the surface of circulating neutrophils, as well as in the serum (Figure S1A-B). Ly6G ligation inhibited neutrophil migration only in integrin-dependent IL-1β peritonitis, whereas no effect could be discerned in the less integrin-dependent *E. coli* peritonitis (Figure 4A-B).

To confirm that reduced neutrophil entry reflected specific inhibition of integrin-mediated migration, we repeated these experiments using our adoptive transfer system. Mice were treated with 1A8, 2A3, and i.p. IL-1β or E. coli, as above. We then transferred a 1:1 mixture of WT neutrophils stained with PKH67 (green) and CD18^-/-^ neutrophils stained with CellVue maroon (far-red) i.v. and quantitated peritoneal cells 3h later by flow cytometry (Figure 4A). WT and CD18^-/-^ neutrophils migrated comparably in *E. coli* peritonitis, and 1A8 treatment had no impact on either population (Figure 4C). By contrast, in IL-1β peritonitis, WT neutrophils showed a migratory advantage over CD18^-/-^ neutrophils, but this difference was less marked in the 1A8- than in the 2A3-treated group (p=0.0137 vs p=0.002) (Figure 4C). Correspondingly, 1A8 altered the ratio of WT to CD18^-/-^ migrated neutrophils only in IL-1β peritonitis (Figure 4D). These results demonstrate that Ly6G ligation selectively impairs β2 integrin-dependent neutrophil migration.

**Figure 4.**
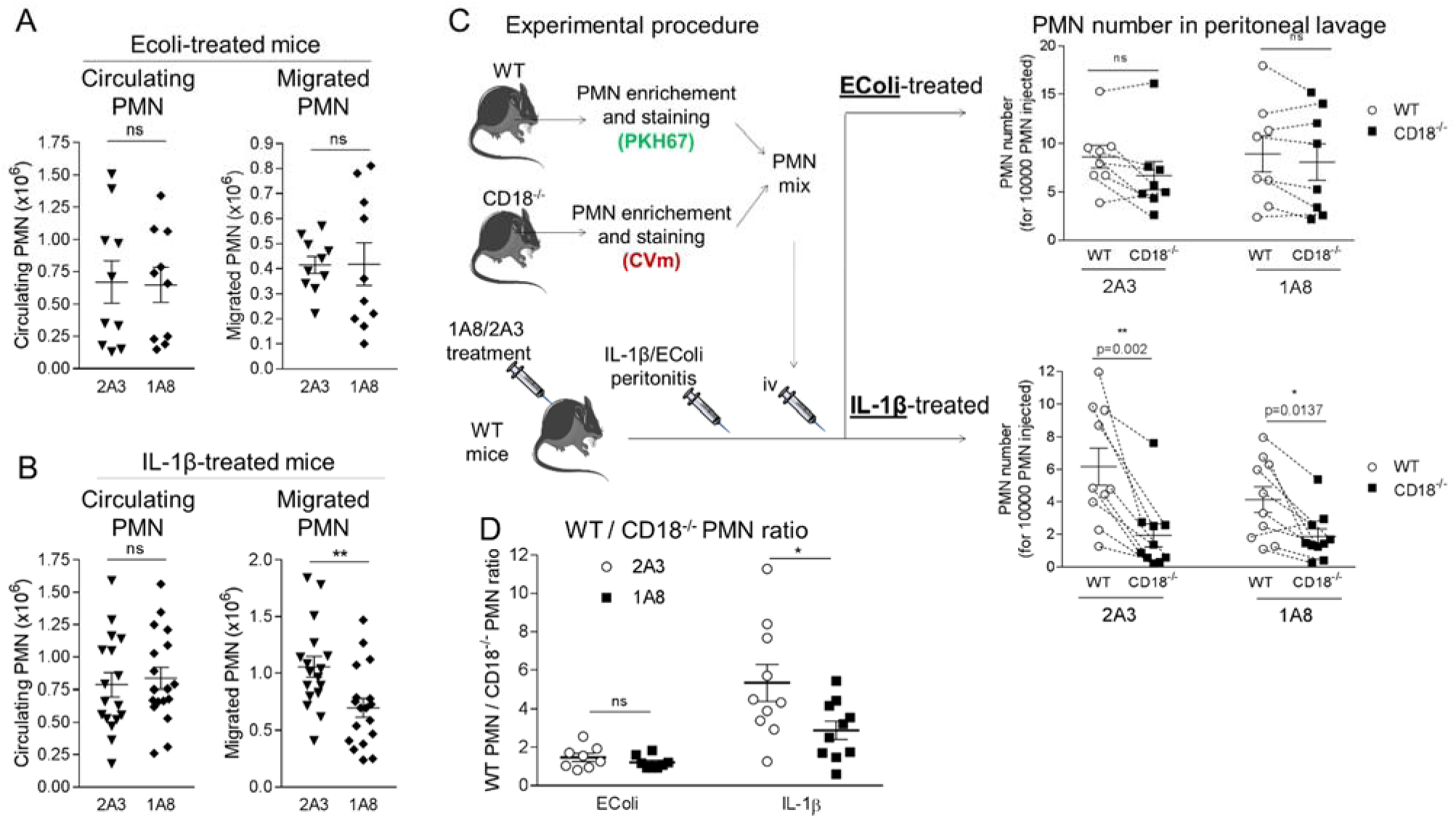
Ly6G ligation selectively impairs β2 integrin-dependent neutrophil migration into peritoneum. **A-B**. Mice are injected i.p. with 5μg of anti-Ly6G (clone 1A8) or isotype control (clone 2A3) at day 0 and day 2. Five days later, mice were injected i.p. with *E. coli* (**A**) or IL-1β (**B**). Circulating (left panels) and migrated (right panels) PMN were enumerated by flow cytometry, n=10-18 mice per group. **C**. Mice injected ip with 1A8 or 2A3 at day 0 and day 2 were treated at day 5 with *E. coli* or IL-1β prior injection of a mix of WT PMN stained with PKH67 (green) and CD18^-/-^ PMN stained with CVm (far-red), as in Figure 3E. Three hours later, WT and CD18^-/-^ PMN in peritoneal lavage were counted by flow cytometry. **D**. Calculation of the ratio of WT PMN / CD18^-/-^ PMN that migrate during peritonitis, n=8 per group in **E**. coli-treated mice and 14 per group in IL-1β-treated mice.

### Migratory blockade is associated with altered interaction between Ly6G and β2 integrins

To understand how Ly6G ligation selectively interrupted integrin-mediated migration, we characterized the association of Ly6G with β2 integrins in our experimental animals. Confocal microscopy of fixed peritoneal neutrophils from E. coli- and IL-1β-treated mice confirmed colocalization of these molecules on the neutrophil surface, greater in *E. coli* peritonitis than in IL-1β peritonitis (Figure 5A). To further quantitate this change in spatial association, we employed fluorescence lifetime imaging microscopy (FLIM). FLIM assesses the proximity between two molecules via image-based quantitation of donor fluorescence lifetime, which decreases upon fluorescence resonance energy transfer (FRET) to an acceptor chromophore. As shown in Figure 5B, the fluorescence lifetime τ_1_ of the donor (anti-CD18) decreased in the presence of 1A8 (anti-Ly6G, acceptor), confirming proximity of CD18 and Ly6G, most prominently in *E. coli* peritonitis. We then tested the effect of 1A8 on this molecular interaction. Fresh bone marrow neutrophils were incubated with 1A8 or 2A3 with or without activation by LTB4; cells were then fixed and imaged by FLIM. In the absence of 1A8, Ly6G and CD18 interacted more closely in activated than in resting neutrophils, as reflected in decreased τ_1_ (Figure 5C). However, 1A8 fully abrogated this change (Figure 5C), suggesting that an altered relationship between these molecules could contribute to the migratory blockade observed *in vivo*.

**Figure 5.**
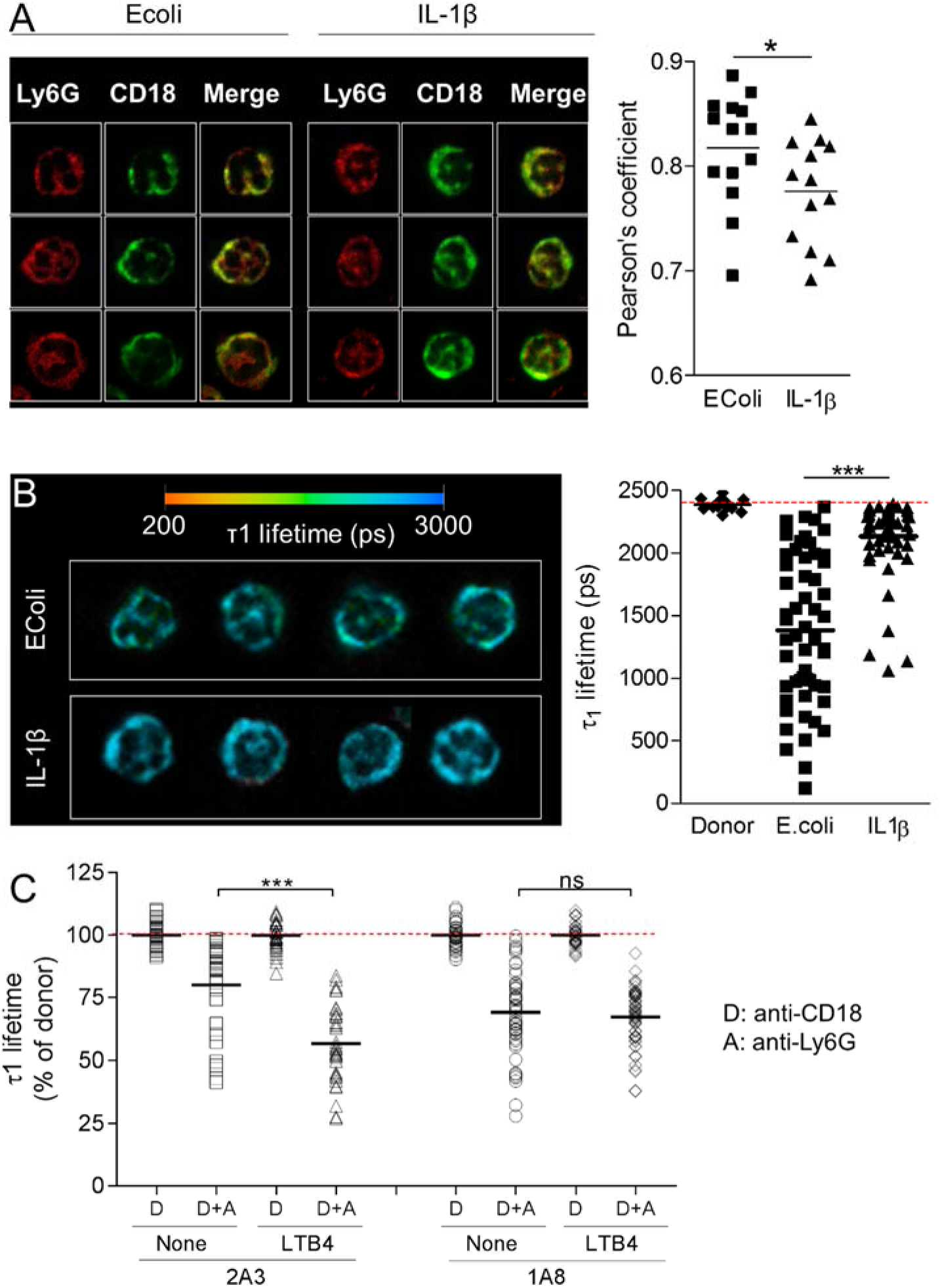
Ly6G ligation modulates the spatial relationship of Ly6G with β2 integrins. A. Mice were treated with either *E. coli* or IL-1β i.p. 3 hours later, cells from peritoneal lavage were fixed and stained with Ly6G (red) and CD18 (green) mAbs. Left, representative photos from confocal microscope. Right, Pearson’s coefficient in PMN from *E. coli* or IL-1β-treated mice. **B**. Peritoneal PMN from *E. coli*- or IL-1β-treated mice were stained with FITC anti-CD18 (donor) and AF594 anti-Ly6G (acceptor) mAbs. Left, representative images from each group displaying interacting τ_1_ lifetime in each pixel on a pseudocolor scale. Right, τ_1_ lifetime in at least 55 different pixels, representative of 3 independent experiments. **C**. PMN were incubated with 10μg/ml 1A8 or 2A3 and stimulated or not with 20nM LTB4. After fixation, cells were stained with anti-CD18 (donor, “D”) without or with anti-Ly6G (acceptor, “D+A”) for FLIM. At least 55 different pixels per conditions were analyzed. Results are expressed as a percentage of τ_1_ lifetime in donor conditions and are representative of 2 independent experiments.

Finally, we imaged neutrophil-endothelial interactions in exteriorized cremaster muscle, a system in which the role of β2 integrins has been intensively characterized.^13,16,28^ Among rolling cells tracked for at least 15 seconds after CXCL1 injection, rates of initial arrest were equal between isotype (32 of 33 cells, 97%) and 1A8 (37 of 38 cells, 97%). However, compared with isotype control, 1A8 markedly shortened the duration of endothelial residence induced by i.v. injection of CXCL1, reflecting impairment of firm endothelial attachment (postadhesion strengthening), a distinct integrin-mediated phase of neutrophil transendothelial migration (Figure 6A and Supplementary Videos 1 and 2).^1-3^ Analysis of the leukocyte rolling flux in postcapillary venules confirmed that the stability of neutrophil firm adhesion was impaired by Ly6G ligation, as return to prestimulation leukocyte rolling flux occurred more rapidly in 1A8 treated mice (Figure 6B). Thus, consistent with the studies in arthritis, pneumonitis, and peritonitis, Ly6G ligation modulated integrin-dependent behavior in neutrophils directly visualized *in vivo*.

**Figure 6.**
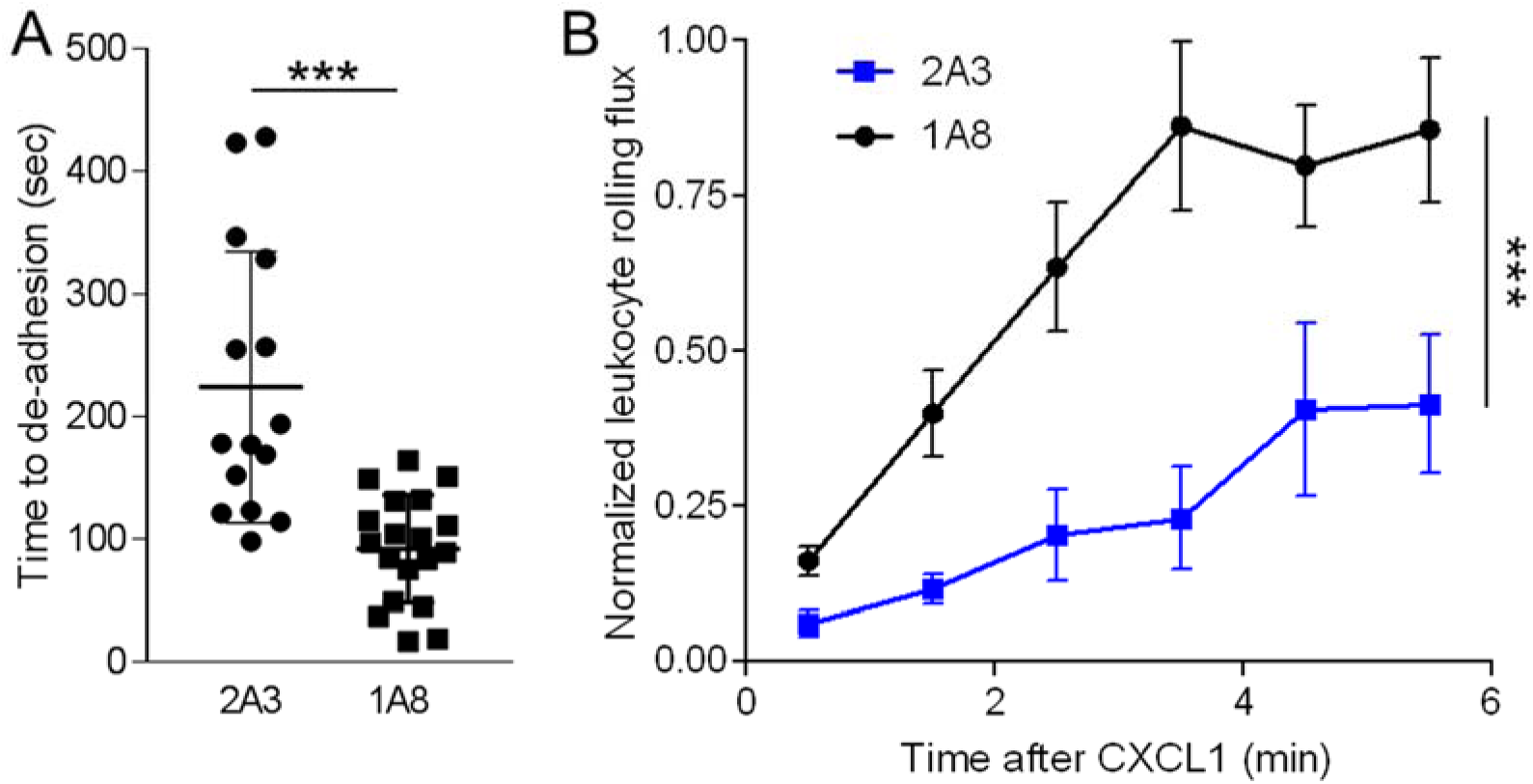
Ly6G ligation impairs neutrophil attachment to endothelium. **A-B**. Mice were treated with 5μg 2A3 or 1A8. Four hours later, leukocyte arrest in postcapillary venules of the cremaster muscle is induced by injection i.a. of 1μg CXCL1. **A**. Duration of attachment of neutrophils on the endothelium in seconds. **B**. Rolling flux was quantified as the number of leukocytes rolling past an arbitrary landmark, counted in one-minute segments and then normalized to the leukocyte rolling flux prior to CXCL1 injection. n = 5 mice per group.

## Discussion

Neutrophil migration into tissues is a complex process. In most contexts, β2 integrins play a key role, mediating firm adhesion as well as diapedesis. Yet studies using genetically-deficient animals, blocking antibodies, and pharmacologic manipulation confirm that the integrin contribution can be surprisingly variable. The principles governing when integrins are required and when not remain to be fully elucidated. The present studies confirm that dependence of neutrophil migration on β2 integrins ranges from largely required (the inflamed joint) to largely dispensable (lung) and can vary with stimulus. We find further that ligation of the ubiquitously-expressed murine neutrophil protein Ly6G selectively antagonizes β2 integrin-mediated migration, helping to define how this intervention alters migration in some contexts but not others.^15,19^

Our data extend a substantial body of experimental literature detailing the role of β2 integrins in neutrophil migration. Neutrophils express the β2 integrins LFA-1 and Mac-1, and in lower abundance complement receptor 4 (CD11c/CD18). The role of LFA-1 and Mac-1 in migration has been studied in many disease models. Where published results diverge, it can be difficult to differentiate organ- and/or stimulus-specificity from factors such as animal strain and microenvironment. Since β2 integrins are expressed on many lineages, and participate in neutrophil functions beyond endothelial adhesion (for example, lineage development, pathogen elimination, apoptosis, and swarming), interpretation of differences between WT and integrin-deficient animals can be challenging.^13,23,25,29,30^ Nevertheless, our observations concord with published data, confirming that the role of β2 integrins in neutrophil migration varies with site and stimulus. Our findings in peritoneum echo those in the hepatic sinusoids, where sterile but not septic inflammation displays integrin dependence.^14^ Collectively, these studies highlight the plasticity of pathways mediating neutrophil egress into tissues.

The present work further extends the understanding of the biology of Ly6G. A small protein of approximately 25 kDa, Ly6G is a member of the LU (Ly6/uPAR, urokinase plasminogen activator receptor) family of proteins and is tethered to the cell surface via a glycosylphosphatidylinositol (GPI) linker.^31^ Ly6G is expressed in murine neutrophils regardless of their localization or activation state.^31-33^ Eosinophils may also express this protein at low levels.^34^ Ly6G exhibits a "three finger fold” motif stabilized by disulfide bonds, believed to create a docking site for other molecules, although no counter-ligand has been identified.^31^ Ly6G is often targeted to deplete, visualize, or sort neutrophils, using either 1A8 or the less specific RB6-8C5, which also recognizes Ly6C and thus marks inflammatory monocytes and dendritic cells.^15,19,32,33,35^ Ly6G-deficient neutrophils appear normal in number and function, with intact migration in LPS-induced peritonitis, experimental autoimmune encephalomyelitis, *Aspergillus fumigatus* lung infection, and skin inflammation ^20^ as well as K/B×N serum transfer arthritis (this study). Nevertheless, ligation of Ly6G with either 1A8 or RB6-8C5 elicits functional consequences that can include activation of intracellular signaling cascades and anaphylaxis-like shock in TNF-or LPS-treated mice.^36-38^ We find that Ly6G ligation alters its spatial association with β2 integrins and impairs firm adhesion to activated endothelium in cremaster muscle. The latter phenotype suggests compromise of postadhesion strengthening, a phenomenon mediated predominantly by LFA-1, whereas Mac-1 mediates adhesion-associated intraluminal crawling.^16,39^

The present observations extend the similarities and contrasts between Ly6G and CD177.^31,40,41^ CD177 is another GPI-anchored member of the LU protein family, and in humans it is expressed predominantly by neutrophils.^42,43^ Like Ly6G, CD177 resides on the cell surface in molecular association with β2-integrins, and its ligation by specific antibodies inhibits neutrophil migration.^44-46^ Intriguingly, adhesion blockade resulting from CD177 ligation with the antibody MEM166 reflects enhanced rather than diminished integrin-mediated adhesion, a “leukadherin”-like immobilization mechanism mediated at least in part through cell activation by signaling via the integrin itself.^45^ Whether Ly6G could engage a similar mechanism in a context-dependent manner, or upon ligation via a different set of epitopes than engaged by 1A8 and RB6-8C5, remains to be determined. The identification of endogenous soluble or cellular counter-ligands for Ly6G and CD177 will be essential to understanding the roles of these receptors in health and disease.

A further parallel between CD177 and Ly6G is the surprising paucity of phenotype from CD177 deficiency. A CD177 allele containing a stop codon acquired from a nearby pseudogene is relatively common, such that 3-5% of the human population lacks CD177 without evident phenotypic consequences.^43,45,47,48^ Mice deficient in *Cd177* are relatively normal, exhibiting only a small delay in neutrophil accumulation in *S. aureus*-infected skin but not in thioglycolate-induced peritonitis.^49^ Ly6G and *CD177* may therefore both participate in redundant or otherwise inconsequential mechanisms. However, their evolutionary preservation, close interactions with surface integrins, and the striking consequences induced by ligation suggest that they have physiological functions yet to be identified.

We recognize strengths and limitations of these studies. Our data reconcile apparently conflicting observations within the larger context of studies into the role of β2 integrins in neutrophil migration. Methodologically, adoptive transfer of WT and CD18^-/-^ neutrophils enabled careful dissection of lineage-specific and cell-intrinsic migratory requirements, overcoming the confounding effects of aberrant neutrophil number and distribution in CD18^-/-^ mice. Given the variability of neutrophil migration, we do not claim that the selectivity of Ly6G ligation for integrin-mediated migration extends to every context, or that it explains the whole divergence in observed findings. For example, while the role of β2 integrins in neutrophil recruitment to intradermal *S. aureus* has not to our knowledge been defined, related studies and clinical experience in patents with CD18 deficiency suggest that some contribution is likely.^12,19,50^ We have not defined the molecular mechanism through which Ly6G ligation impedes integrin-specific migration, including the functional consequences of the disrupted association between Ly6G and CD18 identified by FLIM. Interestingly, we found that Ly6G strongly interacts with CD18 on neutrophil surface after *E. coli* treatment. This observation suggests that the Ly6G/CD18 pathway could be activated by *E. coli* as well. However, in this model of septic inflammation, utilization of β2-independent pathways could explain the absence of inhibition induced by 1A8 treatment. Further study will be required to clarify these pathways in physiological and pathophysiological situations.

While exclusively murine, our findings have therapeutic implications. Global impairment of neutrophil number and function likely carries an unacceptable risk of infection, but selective impairment of integrin-dependent neutrophil migration could potentially block some neutrophil functions while preserving others. In this context, it is especially intriguing that integrins contributed more to neutrophil recruitment in sterile than in septic peritonitis, as previously shown in liver.^14^ If this pattern holds true in other contexts, then interference with integrin-mediated neutrophil migration could attenuate pathogenic neutrophil infiltration while sparing at least a portion of their defensive role.

## Material and methods

### Mice

Wild-type C57BL/6J, and B6.129S7-*Itgb2*^tm2Bay^/J (CD18-/-) mice were obtained from The Jackson Laboratory. C57BL/6-Ly6gtm2621(Cre-tdTomato)Arte ("Catchup” mice, Ly6G-deficient) were obtained as described. ^20^ K/BxN mice expressing the TCR transgene KRN and the MHC class II molecule A^g7^ were generated at a dedicated colony maintained at The Jackson Laboratory. Experiments were approved by the animal care and use committees of the Brigham and Women’s Hospital and Rhode Island Hospital.

### Antibodies and reagents

Anti-CD18 (clone M18/2), -CD11b (M1/70), -Gr1 (RB6-8C5), - Ly6C (HK1.4), -Ly6G (1A8), -CD45 (30-F11), and anti-rat IgG (Poly4054) antibodies were from Biolegend. Counting beads (15μm) were from Polyscience Inc. Endotoxin-free anti-Ly6G (clone 1A8) and isotype control (rat IgG2a, clone 2A3) were from BioXCell. CellVue Maroon, PKH67, recombinant murine IL-1β and LPS 0111:B4 were from Sigma Aldrich. *E. coli* were from ATCC, # 25922. LTB4 was from Cayman Chemicals.

### K/B×N serum transfer model of arthritis

150μl of pooled serum from 9-11 week old K/B×N mice were transferred i.p. on day 0 and day 2. Arthritis was graded using a 0-12 clinical scale (0-3 per paw) as described 15, as well as by caliper of wrists and ankles.

### Pneumonitis model

Mice were anesthetized with Ketamine / Xylazine and IL-1β (50ng) or LPS (1μg) in 30μl PBS were administrated intranasally. After 6 hours, mice were sacrificed, and BAL and lungs were harvested. Lungs were digested with collagenase type 4 (2mg/ml, Worthington Biochemical) and DNAse I (0.1mg/ml, Roche) for 30min at 37C. In some experiments, 5μg 1A8 or 2A3 were injected ip 18 hours before lung inflammation. In other experiments, a mix of WT and CD18^-/-^ PMN, stained with either PKH67+ (green) or CellVue Maroon+ (CVm+, far-red – colors varied across experiments) were injected i.v. 1 hour after lung inflammation.

### Peritonitis model

Mice were injected ip with *E. coli* (10^7^ CFU/mouse) or IL-1β (5μg/mouse) in 200μl PBS. After 3 hours, peritoneum was washed with 5ml cold PBS. In some experiments, 5μg 1A8 or 2A3 were injected i.p., 5 days before and once again 3 days before peritonitis. In other experiments, a mix of WT and CD18^-/-^ PMN, stained with either PKH67+ (green) or CellVue Maroon+ (CVm+, far-red – colors varied across experiments) was injected i.v. 1 hour after peritonitis.

### Neutrophil quantitation

Neutrophils in tissues, in lavages or in the circulation were distinguished by flow cytometry based on Ly6G expression. In experiments involving incubation with 1A8, or when mice were treated with 1A8, neutrophils were distinguish based on the markers Ly6C/Gr-1, CD11b/Gr1 or Ly6C/CD11b. Staining were performed in the presence of 1×10^5^ 15μm counting beads. When experiments required injection of green (PKH67) and far-red (CVm) neutrophils, the number of migrated green and far-red cells in tissues were normalized with the number of green and far-red cells present in the mix, before injection (not shown).

### Ly6G/ CD18 colocalization

Neutrophils from peritoneal lavages were fixed with PFA 4%, stained with anti-Ly6G AF594 and anti-CD18 AF488. Immunofluorescence images were taken by a confocal microscope (C1; Nikon) using a Plan Apochromat 60×/1.40 numerical aperture oil objective (Nikon). Pearson’s coefficients were obtained with ImageJ software

### Ly6G/ CD18 interaction

Peritoneal cells, or neutrophils stimulated or not with 20nM LTB4 for 30min were fixed, permeabilized, and stained for fluorescence resonance energy transfer (FRET) donor-acceptor pair, with AF488 anti-CD18 served as the donor fluorophore and AF594 anti-Ly6G served as the acceptor fluorophore. Protein interactions were determined by fluorescence lifetime imaging microscopy (FLIM) using a time-correlated single-photon counting confocal FLIM system Nikon TiE microscope with a Plan Apochromat VC 60×/1.4 DC N2 objective.^15^

### Mouse cremaster intravital microscopy

These studies were approved by the Lifespan (Rhode Island Hospital) Animal Welfare Committee. Mice were anesthetized with ketamine (100 mg/kg) and xylazine (10 mg/kg) during carotid artery catheterization and exteriorization of the cremaster muscle. Leukocyte rolling and adhesion in postcapillary venules (approximately 25-40 μm in diameter) was imaged by brightfield transillumination using an Olympus BX60 microscope and 20X water-dipping objective (Olympus), and recorded at 30 frames per second on a Chameleon3 color CMOS camera (FLIR Integrated Imaging Solutions). Videos were recorded, starting 1 minute prior to injection of recombinant murine CXCL1 (1 μg) via a carotid artery catheter, and proceeding for 6 minutes after injection.

### Statistics

Unless otherwise stated, comparisons between two paired conditions employed the Wilcoxon matched pairs test; comparisons between two groups of mice employed the Mann-Whitney test. For analysis of WT and CD18^-/-^ neutrophil migration in vivo, the ratio WT/CD18-/- was compared to a hypothetical value of 1, using a one-sample t test. For measurement and scoring of arthritis over time, we used the two-way analysis of variance (ANOVA). For FLIM experiments, data were analyzed with Mann-Whitney test. For analysis of leukocyte rolling flux, data were log-transformed prior to analysis by Student’s t-test of curve-fit parameters. Statistical analyses were performed with GraphPad Prism software. Error bars represent SEM; *p≤0.05, **p≤0.01, ***p≤0.001.

## Supplemental Material

### Supplemental Figure1

**Figure S1.**
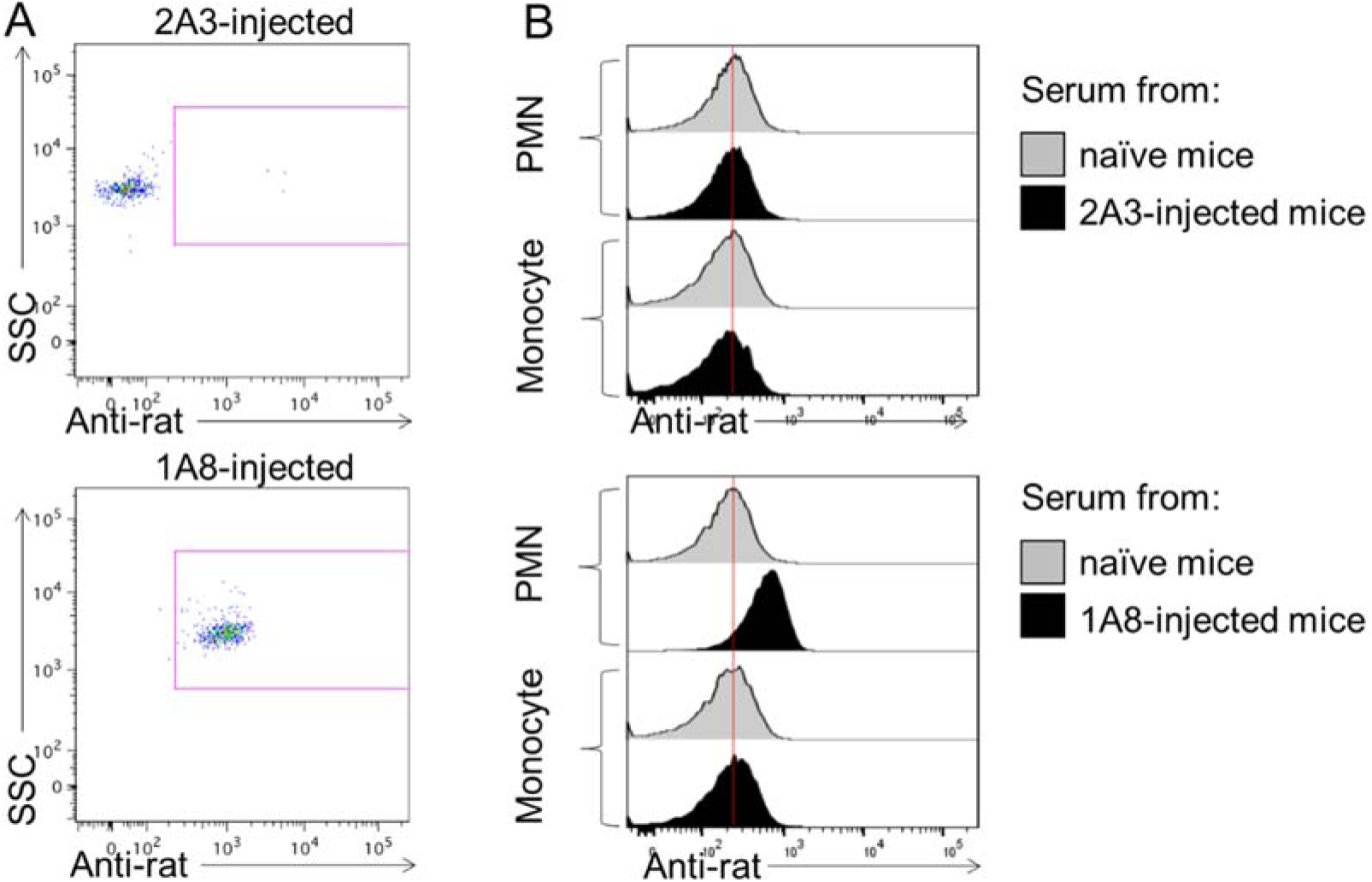
Persistence of anti-Ly6G antibody on circulating neutrophils and in the circulation. Mice were injected i.p. with 5μg of anti-Ly6G mAb (clone 1A8) or isotype control (clone 2A3) at day 0 and day 2. A. Flow cytometry revealed the presence of 1A8 on circulating PMN 5 days after the initial antibody treatment. Representative of at least 3 mice. B. WT bone marrow cells were incubated with diluted serum (1/10 dilution) from 1A8- or 2A3-treated mice, obtained at different time points. The presence of 1A8 or 2A3 on bone marrow cells was revealed with an AF647-labelled anti-rat antibody. After washings, cells were incubated with an anti-Ly6C and anti-CD11b to distinguish PMN (Ly6C^int^/CD11b^hi^) and monocytes (Ly6C^hi^/CD11b^hi^)”

**Supplemental videos:** Two examples of videos obtained with the cremaster intravital microscopy technique. Mice are treated ip with 5μg 2A3 (**video 1**) or 1A8 (**video 2**). Four hours later, leukocyte arrest in postcapillary venules of the cremaster muscle is induced by injection i.a. of 1μg CXCL1. The CXCL1 injection time is 0 minutes 46 seconds in the video 1, and 0 minutes 53 seconds in the video 2.

## ACKNOWLEDGEMENTS

The present study was funded by NIH awards R01 AR065538 and P30 AR070253, and by the Fundación Bechara (P.A.N.). P.C. was supported by a grant from the Arthritis National Research Foundation and a JBC Microgrant from the Joint Biology Consortium (P30 AR070253). P.Y.L. was supported by an Investigator Award from the Rheumatology Research Foundation and NIH T32 AI007512, and a JBC Microgrant from the Joint Biology Consortium (P30 AR070253). E.K. was supported by the American Heart Association (15FTF25080205). E.B. was supported by a Foundation grant from the Canadian Institutes of Health Research (CIHR). C.T.L. was supported by NIH award R35 GM124911. All authors declare no related conflict of interest.

## AUTHOR CONTRIBUTIONS

P.C. conceptualized and designed research, performed research, analyzed data, and wrote the paper. P.Y.L. designed research, performed research, analyzed data, and contributed to writing the paper. E.K. designed research, performed research, and analyzed data. A.B.S. performed research and analyzed data. N.C. performed research and analyzed data. A.P. performed research and analyzed data. M.G. provided critical reagents. R.J.S. supervised research and analyzed data. S.L. supervised research and analyzed data. E.B. supervised research and analyzed data. C.T.L. designed research, performed research, analyzed data, and contributed to writing the paper. P.A.N. conceptualized, designed and supervised research, analyzed data, and wrote the paper.

